# Patient-derived HCC cells recapitulating the transcriptomic landscape of primary HCV-related liver cancer

**DOI:** 10.1101/2024.12.01.626251

**Authors:** Janine Kah, Lisa Staffeldt, Tassilo Volz, Kornelius Schulze, Asmus Heumann, Götz Rövenstrunk, Meike Goebel, Sven Peine, Maura Dandri, Stefan Lüth

## Abstract

**Background:** The hepatocellular carcinoma is one of the leading causes of cancer-related mortality and is characterized by high heterogeneity and subsequently adaptation by developing resistance to current treatments. In this scenario the application of individualized models is crucial to understand the potential of approved therapies. Recently, we established a series of individual cell lines derived from patients who developed HCC on different entities, serving as a platform for individual approaches. In this study, we classified the LC4 cells derived from the center region of a HCC with underlying HIV-HCV co-infection, by using deep analysis on the pathway regulation level.

**Methods:** We employed DEG analysis, followed by pathway analysis to characterize the preservation level of the LC4 cells and the level of adoption. Next, we classify the model, by employing healthy donor samples, commonly used HCC cell lines and global RNAseq data sets.

**Results:** We showed that the LC4 cells reflect significant characteristics of the parental region, including the replication of the immuno-suppressive and the proliferative milieu. The LC4 cells exhibit a metabolic reprogramming characterized by the downregulation of drug-metabolizing CYP enzymes compared to healthy individuals, indicating a transition to alternate metabolic pathways. Moreover, we identified common Biomarkers in the parental tissue, global datasets and the LC4 cells.

**Conclusion:** We showed that the LC4 cell line is applicable as an individual model for pre-clinical testing of treatment regimens in HCC driven research.

## Introduction

Hepatocellular carcinoma (HCC) is one of the most prevalent cancers, ranking as the sixth most common malignancy and the second leading cause of cancer-related mortality [1]. Although the incidence of metabolic steatohepatitis (MASH) of alcohol- or non-alcoholic origin is increasing, chronic viral hepatitis continues to be the predominant cause of liver cancer globally [2, 3]. A significant risk to develop an HCC poses the infection with the hepatitis C virus (HCV) by causing chronic liver inflammation, fibrosis, and cirrhosis [4]. Of note, an additional complication in this setting represents the co-infection with the human immunodeficiency virus (HIV). Whereby individuals co-infected with HIV and HCV exhibit expedited progression to liver cirrhosis and hepatocellular carcinoma facilitate by the synergistic detrimental effects of both viruses [5, 6]. It has been shown that these individuals will more likely develop resistance during therapy.

Despite recently invented checkpoint inhibitors, the 5-year survival rate remain poor [7, 8]. For late-stage HCC patients, the primary goal is therefore prolonging survival while maintaining quality of life. In recent years, immunotherapy has emerged as a promising area, driven by advances in understanding tumor and tumor microenvironment (TME) interactions, as well as the clinical success of immune checkpoint blockade [9]. Here, first-line therapy regimes Atezolizumab and Bevacizumab in combination with antibodies blocking the PD-1/PDL-1 pathway resulting in an increase immune response [10, 11][12]. However, in fact 80-85% of the patients either do not respond or develop resistance during therapy.

The key challenges remain, and lie in elucidating mechanisms, which lead to resistance and efficacy, in identifying novel therapeutic targets, and in developing personalized treatment regimens. To tackle these challenges, commonly used immortalized HCC cell lines, like HepG2, Huh7, and Hep3B have significant limitations in accurately reflecting the complexity of HCC, especially in the aspects of chemoresistance and the effectivity of targeted therapies [13]. Here the incorporation of individual patient-derived cell culture systems has become invaluable for exploring these challenges, by offer more physiologically relevant models, better reflecting tumor heterogeneity and the TME [13, 14]. These models are crucial for studying the molecular pathways driving HCC progression and therapy responses. The use of patient-derived cell lines from diverse etiologies, particularly those with underlying chronical viral infections and comorbidities is necessary to mimic specific in-vivo conditions, thereby enhancing the translational potential of preclinical findings to clinical applications [15]. Building on this, there is a critical need for individual models that account for variability in tumor biology and treatment responses.

In this study, we present a comprehensive analysis of the LC4 cell line, derived from a patient co-infected with HIV and HCV, who later developed HCC. The patient was typed as HLA-A 02:01, 03:01; HLA-B 15:01, 57:01; and HLA-C 01:02, 06:02 positive, with undetectable HIV and HCV titers at the time of resection due to treatment with Ribavirin and Biktarvy. Our study incorporated RNA-based real-time assays, histological and cytometric protein profiling, and RNA sequencing to identify differentially expressed genes (DEGs), pathway alterations, and biomarkers, to classify the LC4 cells in the HCC landscape. Our key findings show that LC4 cells exhibited distinct characteristics reflecting the parental tissue, including the highly proliferative and immunosuppressive character displayed by the upregulation of genes in the PD-L1/VEGF pathway and the downregulation of B2M, IKZF1, and CDKN2A, accompanied by CDK4 overexpression.

## Methods

### Human sample collection

Human HCC-derived samples were collected and ethically approved as described previously [16]. The collective of healthy primary hepatocytes were obtained after isolation of 4 donor liver tissues which were not used for transplantation: TM14 (female, 52years), TM47 (female, 11 month), TM46 (male, 2month), TM52 (female, 30years). The study was approved by the Ethical Review Committee of the Ärztekammer Hamburg (WF-021/11). Handling of the human material was performed in accordance with national guidelines and the 1975 Declaration of Helsinki [17].

### HLA Typing

DNA has been extracted from patient blood cell samples for HLA Class I typing using the DNA Isolation Kit (Wizard HMW DNA Extraction Kit from Promega, GER). DNA was used for Luminex-based high definition LABType rSSO typing (One Lambda, Canoga Park, CA) of the HLA loci HLA-A, HLA-B and HLA-C. LabType SSO is a reverse SSO DNA typing method (rSSO) with sequence specific Oligonucleotide probes that specifically bind to homologous sequence sections of certain HLA alleles. Genomic DNA (5-60ng/µl) was PCR-amplified using locus-specific biotinylated-primers (Exons 2,4 and 5 for HLA-A and -B; exons 2,4,5,6 and 7 for HLA-C) and Amplitaq Polymerase (One Lambda, Canoga Park, CA), resulting in biotinylated-amplicons. The presence of biotin in each amplicon was detectable using R-Phycoerythrin-conjugated Streptavidin (SAPE). The PCR product was denatured and hybridized to complementary DNA probes bound to fluorescently coded beads. Samples were measured by a flow analyzer (FlexMap 3D®, Luminex®, Austin, TX) identifying the fluorescent intensity of phycoerythrin (PE) on each bead. HLA Fusion^TM^ program Version 4.6.1 (One Lambda, Canoga Park, CA) was used to analyze the data.

### HCC cell culture

HCC cell lines were maintained as described previously [18, 19]. For 3D conditions, different amounts of LC4 cells were seeded on BIOFLOAT™ cell culture plates in DMEM containing L-glutamine and glucose, supplemented with 1% P/S, 10% Gibco fetal bovine serum (FBS; all from Thermo Fisher Scientific, Waltham, USA) and were obtained for 3 weeks to follow the spheroid formation. Spheroids were used for RNA Isolation after 24 days of formation when dense and packed spheroids were obtained, as shown in **Supplementary Figure 1**. For visualization, LC4 cells stably transduced with the vector LeGO-iG2-Puro+-Luc2 (3rd generation HIV1-derived self-inactivating vector) were used. Stable transduction was performed as described previously [16]. For 2D conditions, LC4 were seeded as described previously [16] and used for RNA isolation after forming a monolayer of over 90% density.

### Isolation of Oligonucleotides

RNA was extracted from human liver specimens using the RNeasy Mini RNA purification kit (Qiagen) [20]. RNA was extracted from seeded cells with or without treatment after indicated time points using the RNeasy RNA Micro purification kit (Qiagen).

### Measurement of gene expression level using TaqMan-based PCR

For measurement of gene-expression, 2-step PCR was performed. Therefore, complementary DNA (cDNA) synthesis was conducted by using MMLV Reverse Transcriptase 1st-Strand cDNA Synthesis Kit (Lucigen, Middleton, Wisconsin, USA) to synthesize RNA complementary DNA, according to the manufacturer’s instructions. Human-specific primers from the TaqMan Gene Expression Assay System, listed in **Supplementary Table 1**, were used to determine gene expression levels (Life Technologies, Carlsbad, California, USA). Samples were analyzed using the Quant Studio 7 Real-Time PCR System (Life Technologies, Carlsbad, California, USA). The mean of the human housekeeping genes, glyceraldehyde-3-phosphate dehydrogenase (GAPDH) and ribosomal protein L0 (RPL0) was used to normalize human gene expression levels.

### Flow cytometry

LC4 cells line LC4 were characterized using the monoclonal antibodies anti-CD68-APC700 and anti-CD45-BV510. Cells were stained as described previously, and measurement was carried out on the BD FACSymphony™ A3 Cell Analyzer from BD Biosciences (Heidelberg, GER).

### Protein analysis by immunofluorescence

For histological characterization, cultured cells and tissue slides were processed as described previously [16, 21]. Antibodies used in the study are listed in **Supplementary Table 2.** Stained cells and tissue slides were analyzed by fluorescence microscopy (BZ-9000 and BX-780, Keyence, Osaka, Japan). Captures were automatically generated with the indicated magnification and consistent exposure for the same staining.

### RNA Isolation and Sequencing

Total RNA was extracted from primary hepatocyte samples, primary HCC cell line LC4 and immortalized HCC cell lines (HepG2, Huh7, and Hep3B) using the RNeasy Mini Kit (Qiagen), following the manufacturer’s protocol. The quality and quantity of RNA were assessed using a NanoDrop spectrophotometer (Thermo Fisher Scientific) and Agilent Bioanalyzer 2100 (Agilent Technologies). High-quality RNA samples with an RNA Integrity Number (RIN) greater than 7.0 were selected for sequencing.

### Library Preparation and RNA Sequencing

RNA sequencing libraries were prepared using the TruSeq Stranded mRNA Library Prep Kit (Illumina) according to the manufacturer’s instructions. Briefly, mRNA was purified from total RNA using poly-T oligo-attached magnetic beads and fragmented into small pieces. First-strand cDNA was synthesized using random hexamer primers and reverse transcriptase. This was followed by second-strand cDNA synthesis, end repair, A-tailing, adapter ligation, and PCR amplification to enrich the cDNA fragments. The libraries were quantified using a Qubit fluorometer (Thermo Fisher Scientific) and assessed for size distribution using the Agilent Bioanalyzer 2100. The libraries were then sequenced on the Illumina HiSeq 2500 platform, generating 150 bp paired-end reads. Technical replicates were used for RNAseq for each sample (n=2).

### Data Processing and Differential Expression Analysis

Raw sequencing reads (n = 2 per sample) were prepared as R input data sets and further processed with R [22]. Input was trimmed using Trimmomatic [23]. Clean reads were aligned to the human reference genome (GRCh38) using STAR aligner [24]. The resulting BAM files were processed with featureCounts to generate read count matrices [25]. Differential expression analysis was performed using the DESeq2 package in R [26]. Genes with an adjusted p-value (Benjamini-Hochberg correction) of less than 0.05 and a log2 fold change (log2FC) greater than 2 or less than -2 were considered significantly differentially expressed.

### Data Processing and Visualization

DEGs were used for volcano plot generation, processing comparison analysis in IPA for pathway comparison and in silico analysis of molecules as biomarkers or drug targets in IPA. Results from IPA comparisons were used for bubble plots, z-score based heatmaps and correlation heatmap. The data processing, comparison, and illustration of correlation, pathway comparisons were described in detail in the **Supplementary methods** [27–30].

## Results

### The parental tumor regions are classified as immunogenic margin and an immunosuppressive core

First, we conducted a detailed analysis of spatial heterogeneity of the parental HCC tissue comparing the core and margin region by using expression profiling and immunofluorescence staining. As shown in **Figure 1A** the hepatocyte-specific genes like CLRN3, AADAC, and ALB were decreased in the core region, indicating the transformation from healthy hepatocytes to tumorigenic cells. Conversely, genes such as CD44, GLUT1 and HIF1a were elevated in the margin, suggesting stemness, metabolic adaptation, and hypoxic conditions linked to aggressive tumor biology and potential invasiveness. As shown in Figure 1A and B the proliferation marker Ki67 was upregulated in the core region compared to the margin, which shows signals of hypoxic adaptation and invasion. The protein levels shown in Figure 1B further illustrate these spatial differences, with CD44 and Vimentin (VIM) being more prominent in the margin highlighting the epithelial-mesenchymal transition (EMT) at the invasive front. CD68, a macrophage marker, is present in both the margin and core, indicating tumor-associated macrophage (TAM) infiltration throughout the tumor. However, the roles of these macrophages may vary in the margin, as they promote inflammation and tissue remodeling, while in the core, they contribute to immune evasion through elevated PD-L1 expression. The presented CD3-positive T-cell infiltration in the margin points to a more active immune response, whereas higher PD-L1 expression in the core refers to a consistent immune evasion mechanism.

**Figure 1:**
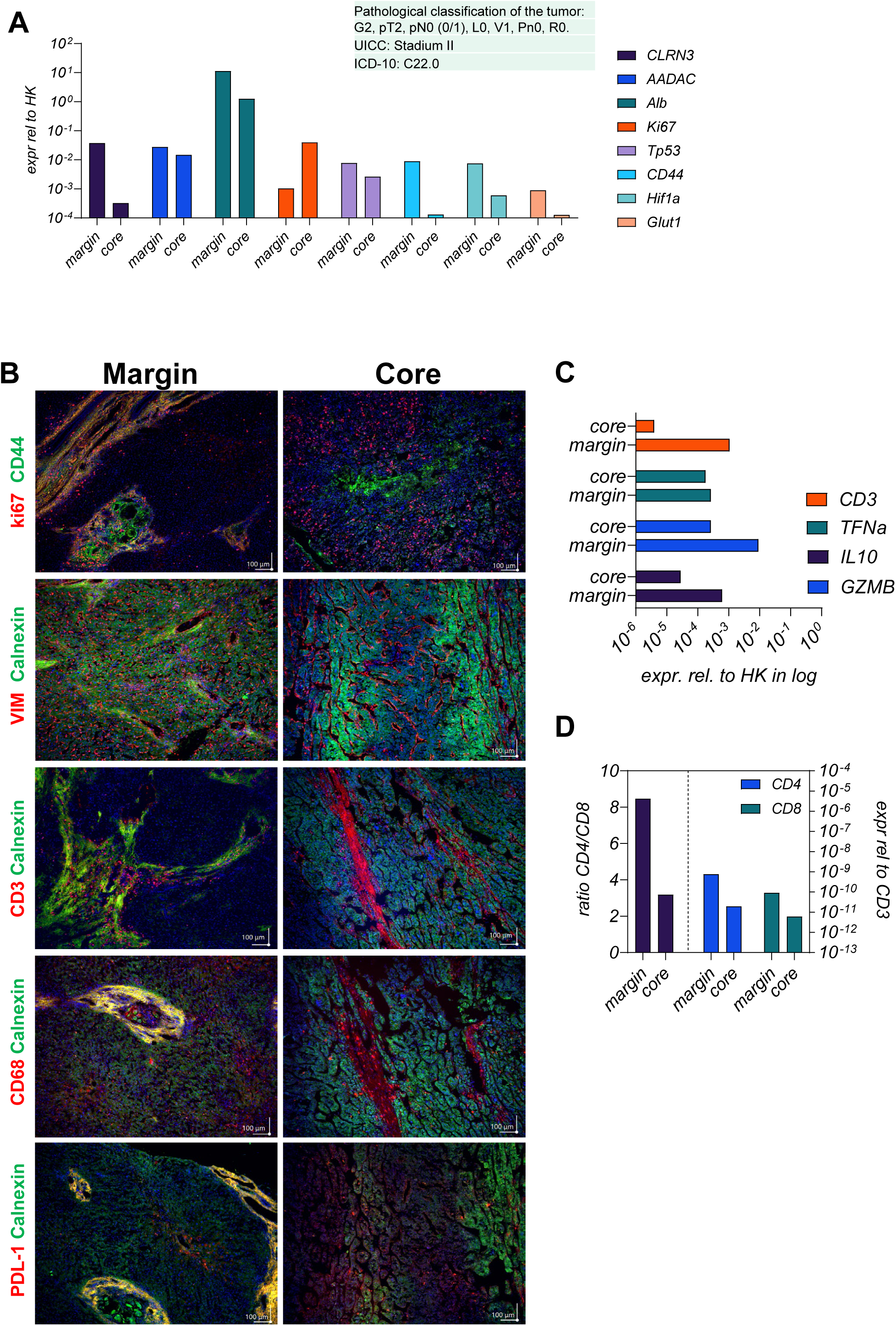
Analysis of gene and protein expression levels in patient-derived tumor tissue regions. **Figure 1A** displays mRNA expression levels of indicated genes resented in the margin and core region of the tumor. Gene expression was normalized relative to HK gene expression (GAPDH and RPL30) and represented on a log10 scale. The analyzed genes include CLRN3, AADAC, Alb, Ki67, Tp53, CD44, Hif1a, and Glut1. The tumor is classified according to pathological criteria: G2, pT2, pN0 (0/1), L0, V1, Pn0, R0. UICC: Stage II, ICD-10: C22.0. **Figure 1B** represents results in 20fold magnification from immunofluorescence staining of the resected tumor tissue comparing the margin and core region. The top row displays the localization of Ki67 (red) and CD44 (green) in both the margin and core, revealing differences in proliferative activity and stemness between these areas. Subsequent rows depict staining of VIM (red) and Calnexin (green), CD3 (red) and Calnexin (green), CD68 (red) and Calnexin (green), and PDL-1 (red) and Calnexin (green). Scale bars represent 100 µm distance. **Figure 1C** presents mRNA expression levels of immune-related genes, including CD3, TNFα, IL10, and GZMB, in the HCC4 tumor core and margin. Expression levels are normalized relative to houser keeper RPL0 gene expression and shown on a logarithmic scale. This comparison illustrates variations in immune response gene expression between the core and margin of the tumor, suggesting differential immune activity within the tumor microenvironment. **Figure 1D** shows the ratio of CD4/CD8 T cells and the expression levels of CD4 and CD8 mRNA in the HCC4 tumor margin and core relative to the T cell lineage marker CD3. The data are presented on a logarithmic scale, highlighting the differences in T cell populations and their gene expression between the margin and core regions of the tumor.

The higher occurrence of CD3 positive T cells in the margin, detected by gene expression levels shown in **Figure 1C** and supported by the protein level shown in **Figure 1B**, was in line with the higher expression of the immunosuppressive cytokine IL10 in the core region. The detection of elevated TNFα in the margin revers to ongoing inflammation at the tumor border. **Figure 1D** shows a higher CD4/CD8 ratio in the margin, indicating a strong helper T-cell response. **Figure 2A** examines the spatial distribution of the HCV core protein (Genotype 1b) and the co-localization of CD3-positive T-cells. As shown in the upper panel, HCV core protein co-localizes with Calnexin, a hepatocyte marker, exclusively in the core of the parental tissue. Despite form the fact, that CD3-positive T-cells were most prominent in the margin, confirming once again the immunosuppressive environment for the core, we found a clear interaction of CD3-positive T-cells with HCV core GT1b positive cells, as shown in the higher magnification lower panel of the **Figure 2A**. The distinct distribution of CD3-positive T-cells between the margin and core draw the spatial heterogeneity of the tumor, where the core, harboring viral proteins, is more adept at suppressing immune infiltration, while the margin maintains stronger immune activity.

**Figure 2:**
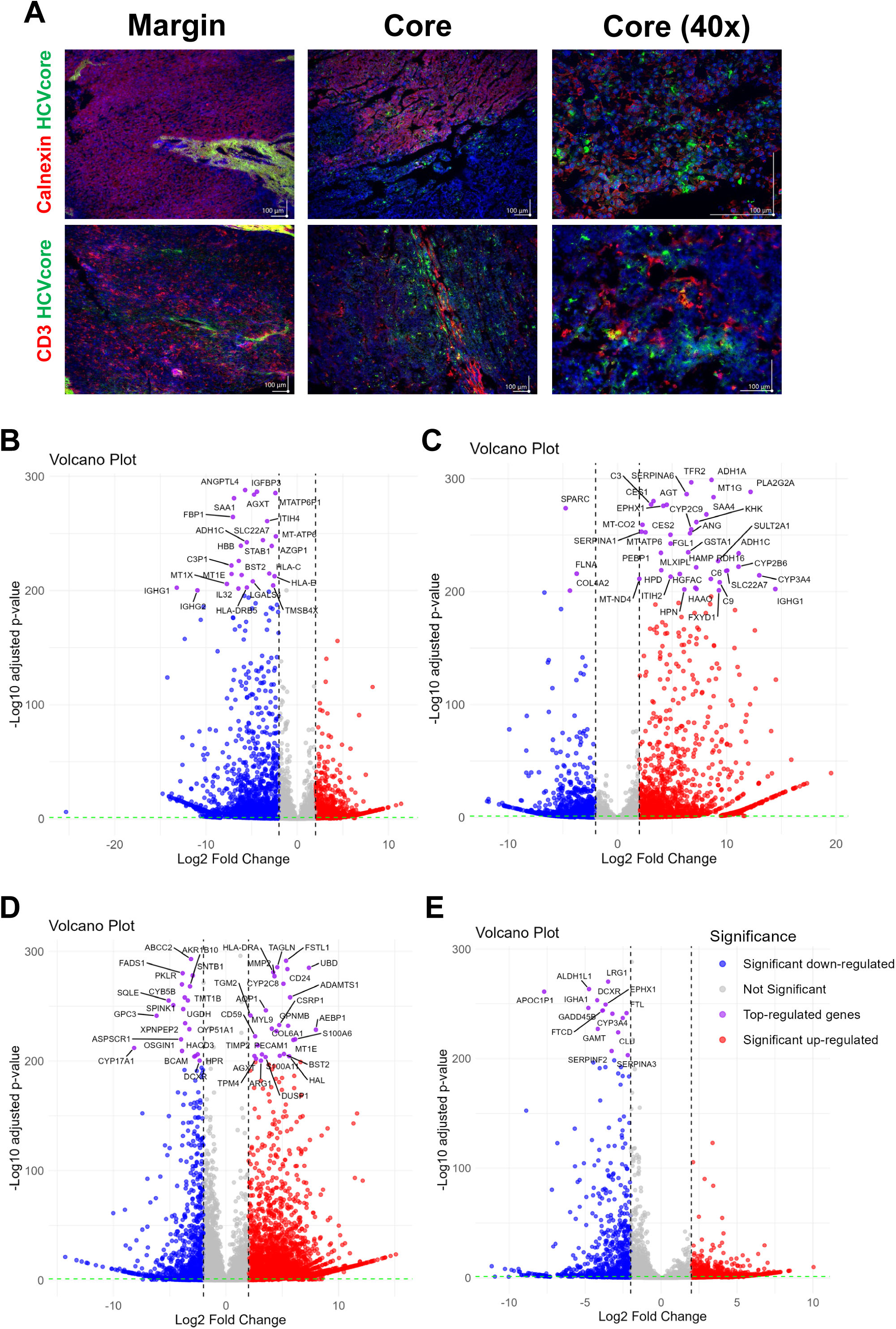
Comparative HCVcore protein occurrence and RNASeq based analysis of tumor regions and conditions. **Figure 2A** presents immunofluorescence staining of HCC tumor tissue, comparing the margin and core regions. The top row shows the localization of Calnexin (red), HCV core (green), and their overlay in both the tumor margin and core. The middle row displays staining for CD3 (red) and HCV core (green), highlighting the immune cell infiltration in relation to viral presence within the tumor. Scale bars represent 100 µm distance. **Figures 2B-E** presenting volcano plots of differential gene expression analysis calculated as followed: B) tumor core tissue against margin tissue, C) tumor core isolate against the margin tissue, D) tumor core tissue against healthy hepatocytes and E) tumor margin tissue against healthy hepatocytes. The x-axis represents the log2 fold change in gene expression, while the y-axis shows the -log10 adjusted p-value, indicating the statistical significance of these changes. Genes significantly downregulated are highlighted in blue, those significantly upregulated in red, and non-significant changes are in gray. The top-regulated genes (p-value <200) are labeled and represented in purple.

In **Figure 2B** we provided a comprehensive view of the differential gene expression (DEG) between the core and margin. In line with our previous findings, genes involved in antigen presentation, such as HLA-B, HLA-C, and HLA-DRB5, are significantly higher expressed in the margin, reflecting the enhanced immune recognition at the periphery compared to the center. The downregulation of metabolic genes and the upregulation of IGFBP3 in the core shows a shift toward proliferation and immune evasion, as shown before. The lower infiltration of CD3-positive T-cells in the core aligns with these expression patterns, further supporting that the tumor core adopts immune-suppressive strategies. Taken together, the tumor tissue from the patient reflects the common picture of most HCCs, whereby the margin is more inflammatory and invasive, while the core exhibits higher proliferation and immune evasion.

For the generation of the LC4 cell line, we employed isolated cells from the core region. Based on this, we aimed to classify the preservation of core region specific characteristics in the LC4 cell line, as environmental changes occur after cultivation. The DEG comparison reported in **Figure 2C** shows a clear shift in the genetic composition due to isolation procedure. In the tumor core, cells face hypoxic conditions, limited nutrient supply, and immune suppression, which keep certain genes downregulated, as shown in **Figure 2B**. After isolation procedure, the core derived cells have been exposed to a more favorable environment with improved oxygen and nutrients, whereby stress responses, metabolic reprogramming, and activation of pathways associated with survival, proliferation, and invasion were triggered. Additionally, isolation simulates wound-healing or tissue-regeneration responses and activating genes involved in extracellular matrix remodeling and immune modulation as mentioned in **Figure 2C**. As in **Figure 2D** and **E** we summarized the transcriptomic status of the core (**D**) and the margin (**E**) in the context of healthy hepatocytes, we found the margin as close-to-healthy region, with only some genes downregulated involved in inflammation and metabolic pathways. On the other hand, the core region represents the same profile as described before with highly significant upregulation of genes involved in cell migration, cytoskeletal restructuring, and immune modulation. In the core, we found GPC3 downregulated, while CD24 was upregulated. Genes like TPM4, TGM2, and S100A11 were significantly upregulated in the core, reflecting enhanced cell migration and invasiveness, consistent with the findings in **Figure 1**. The downregulation of lipid metabolism-related genes like CYP17A1 and SQLE suggests the core reconfigures its metabolic pathways to support aggressive tumor behavior, focusing more on structural adaptation and motility. While showing signs of tumor adaptation, the margin retained normal tissue characteristics, with downregulation of genes involved in iron accumulation, detoxification, and immune response (FTL, SERPINA3, IGHA1). This aligns with earlier observations that the margin shows a less aggressive, more regulated state due to the presence of an active immune response.

### Generated LC4 cell line positively correlates with the parental HCC core region in pathway analysis and shows high similarity to liver cancer related disease

To investigate the preservation of the parental characteristics and to classify patient-derived LC4 cells, we performed DEG analysis from LC4 cells in 2D and 3D culture condition to healthy hepatocytes, to the parental tissues (core and margin) and to the cell isolate derived from the HCC core region, which was used for the generation of the LC4 cell line. Therefore, we performed DEG combinations of all specimens and used the results to acquire the differential affected canonical and toxicity pathways. We next extracted the associated z-scores form the comparison from the Pathway finder software (IPA) and calculated the pairwise Pearson correlation, plotted in **Figure 3A**, whereby the right site represents the canonical and the left site the toxicity pathways. The correlation heatmap matrix visualizes the strength and direction of correlations between the different specimen comparisons, as mentioned in the legend. When correlating the HCC core tissue compared against the margin (X3) or the healthy hepatocytes (X27), we found a positive correlation in both canonical and toxicity pathways. The matrix also clearly shows a positive correlation between both pathway regulations detected in in the LC4 cells when compared to healthy hepatocytes (X31, X32) or the tumor margin tissue (X53, X54) with the HCC core tissue when compared to the tumor margin tissue (X3). When comparing the HCC core tissue against the healthy hepatocytes (X27), a positive correlation was detected to LC4 cells (2D and 3D) compared against the HCC core (X48, X49) and margin (X53, X54) to the same intent. As expected, the isolated HCC core cells when compared to healthy hepatocytes (X30) showed a negative correlation to both LC4 cells (X31, X32) and the parental tissue (X27). This negative correlation was even stronger in the toxicity pathways. Interestingly, the correlation of the canonical pathways differs from the toxicity pathways to some intent, especially, when correlating the parental margin and core both compared to healthy hepatocytes (X26, X27). Here both regions showed a positive correlation in the canonical Pathways, but a negative correlation in the toxicity. As expected, LC4 cells showed mostly positive correlation between the comparisons against healthy hepatocytes, core tissue, the margin and a weak or negative correlation when compared with the isolated cell fraction (X50, X51). Taken together, the LC4 cell line in 2D and 3D conditions were able to reflect the canonical and toxicity pathways regulation as detected in the parental core tissue. To underline these findings, we depicted the underlying volcano plots of the DEGs for X53 (**B**) and X54 (**C**) in Figure **3**. Here the differential gene expression shown in **Figure 3C and 3C** reflects the immunosuppressive character of the parental tumor core region, by downregulation of genes involved in immune response, inflammation, and extracellular matrix remodeling. To classify the generated cell lines in a global setting, we employed the IPA analysis match function. Here we filtered the for datasets and analysis from human liver and set the overall z-score to 10. The highest similarities have been plotted in **Figure 3D and E**, whereby the highest number of studies which show similarity to our analysis was in the HCC in both conditions. This results underline the previously shown results and helps to classify the LC4 cell line as HCC representative cell line.

**Figure 3:**
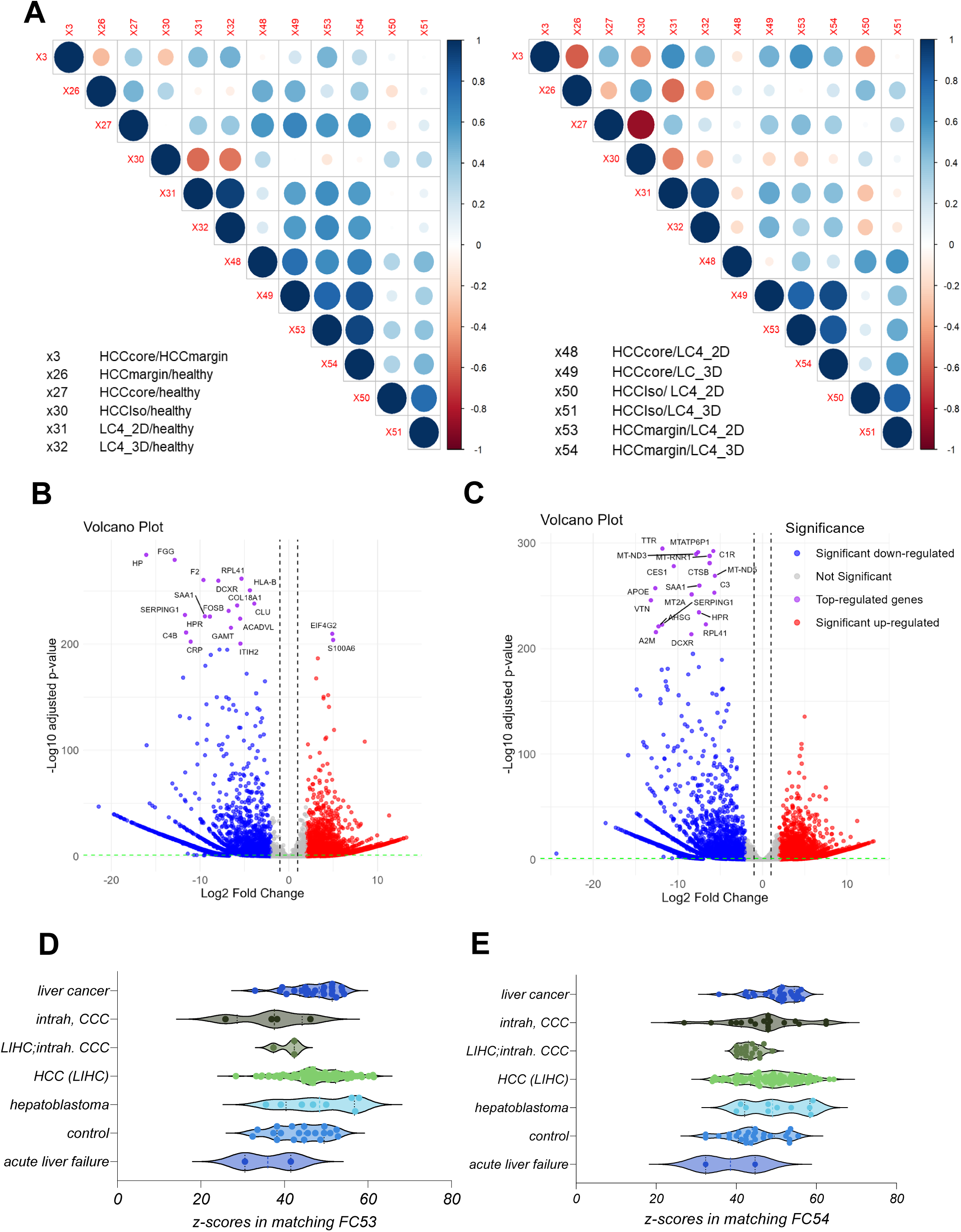
Z-score based correlation analysis of patient-derived specimens and DEG presentation of generated cell line LC4 against tumor regions. **Figure 3A** represents a correlation matrix displaying pairwise Pearson correlations between datasets of calculated z-scores of canonical pathways (right panel) and toxicity pathways (left panel) for the indicated DEG comparisons. Blue and red circles indicate the strength and direction of the correlation, with blue representing positive correlations and red representing negative correlations. The size of the circle corresponds to the magnitude of the correlation coefficient, ranging from -1 to 1. **Figures 3B and C** represent volcano plots of DEG comparisons between generated cell line LC4 in 2D (B) and 3D (C) conditions against the margin tissue. The volcano plots illustrate the log2 fold change in gene expression (x-axis) versus the -log10 adjusted p-value (y-axis). Each point represents a gene, with the x-axis showing the log2 fold change in expression levels between the margin and centrum and the y-axis showing the -log10 adjusted p-value. Blue points indicate significantly down-regulated genes (adjusted p-value < 0.05), red points indicate up-regulated considerably genes, and purple points highlight the top-regulated genes based on fold change. Grey points represent genes with non-significant changes, defined by adjusted p-value > 0.05 and log2 fold change between 2, -2. The top-regulated genes (p-value <200) are labeled and represented in purple. In **Figure 3D and E**, the highest matches, indicated by high z-scores, from the analysis match function in IPA was visualized form the different disease subtypes. In D, we compared the analysis between LC4_2D cells and the margin tissue (FC53) and in E, we show the global comparison match with the 3D condition (FC54) compared to margin tissue from the origin. Every dot represents one single analysis of the global dataset from IPA. For matching, the analysis was reduced to liver, human and the overall z-scores higher or lower 10. In the graphs D and E, all highest similarities were plotted and group as in IPA.

### LC4 Cell Lines Retain Core Tumor Traits but Exhibit Enhanced Proliferation and Reduced Stress-Related Pathways Compared to Parental HCC Core

Based on the correlation matrix, shown in Figure 3A, we focused on a deep pathway activity analysis of parental HCC core tissue (FC3) with LC4 cell lines derived from it, grown in both 2D (FC53) and 3D (FC54) cultures. As shown in Figure 4A, LC4 cells in both 2D and 3D culture display similarities in metabolic pathways to the parental HCC core tissue. However, LC4 cells show enhanced activity in pathways related to transcription, protein processing, and translation, as shown in Figure 4B. This shows, that LC4 cells and particularly in 3D culture, exhibit a partially higher biosynthetic demand, reflecting the adaptation to in-vitro conditions that differs from the in-vivo environment. As summarized in Figure 4B higher activation of metabolic pathways were detected in LC4 cells under different conditions, indicating that LC4 cells shifting their metabolic activities from the stress-adapted metabolic profile in the core. As shown in the Figure 4C, LC4 cells displayed elevated activity in pathways associated with viral infections such as viral hepatitis (8), inflammation of the liver (7), and hepatitis B infection (1). This shows that LC4 cells recapitulate aspects of immune and viral stress pathways, possibly due to changes in the microenvironment under in-vitro conditions. As expected, the liver tumor pathway (10) is more activated in the parental HCC core than in the LC4 cells.

**Figure 4:**
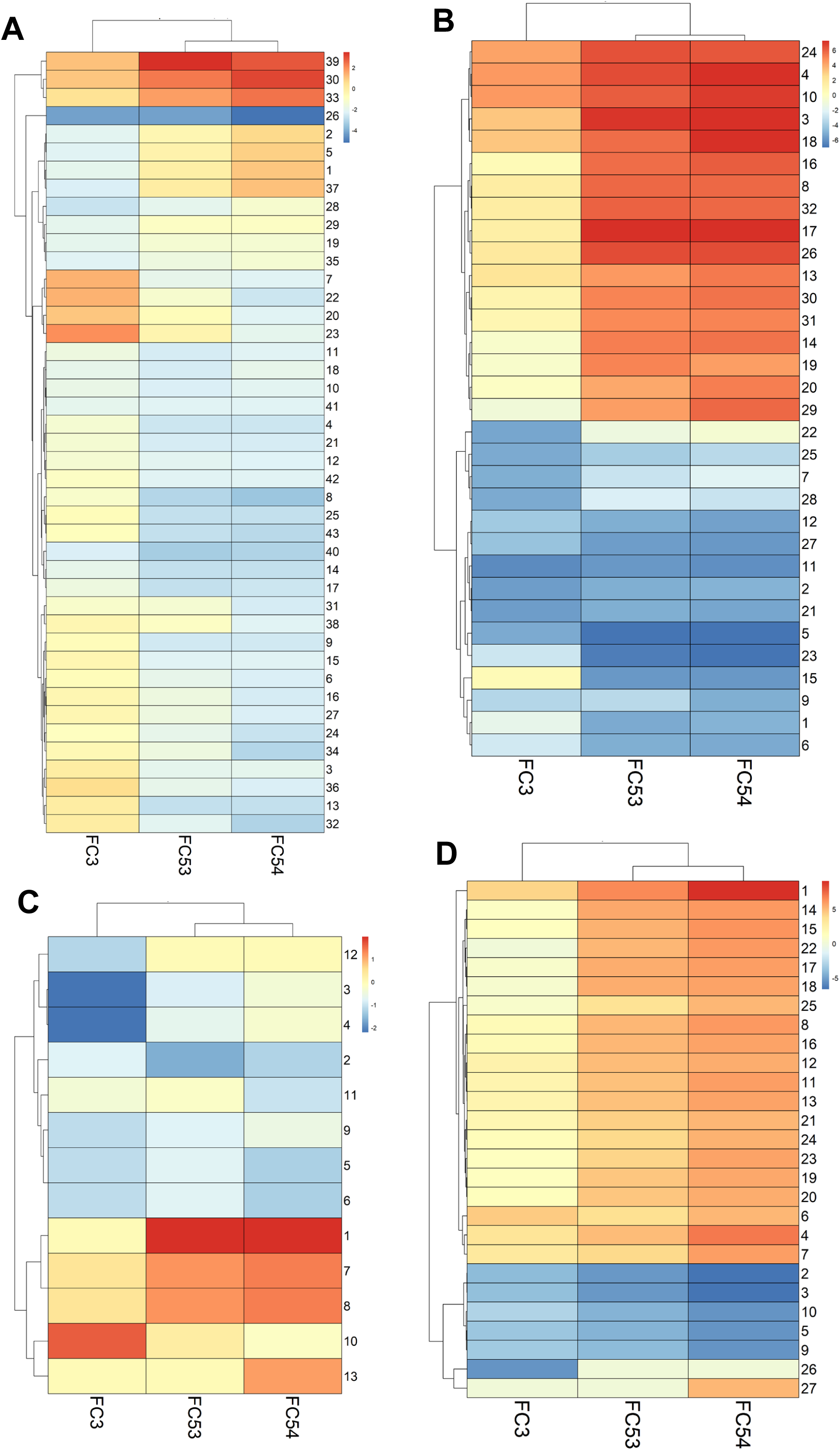
Comparison of relevant canonical pathways, disease & biofunctions, toxicity functions in patient-derived specimens. **In Figure 4** pathway comparison was performed form DEG analysis, calculated against the tumor margin expression for the experimental conditions: core tumor tissue (Analysis 3), LC4 cells in 2D culture (Analysis 53), and LC4 cells in 3D culture (Analysis 54). Z-scores > -5 and < 5 with log_10_p-values > 1,3 have been compared. The heatmaps showing the z-scores of pathway activation, with red indicating upregulation and blue indicating downregulation. In panel **A** metabolic canonical pathways and in panel **B** signaling canonical pathways are shown for the three comparisons. In panel **C**, the toxicity functions and in panel **D**, diseases and biofunctions were compared, between the generated cell line LC4 and the core region of the parental tumor. For comparative illustration pathways were numbered using pheatmap package in Rstudio and decoded in the **table 1-4**.

LC4 cells, particularly in 3D culture, demonstrate higher predicted activity in proliferation of hepatic stellate cells (4) and liver cells (3) compared to the HCC core, due to the absence of growth-inhibitory signals or stress factors present in the tumor microenvironment. Additionally, in the Figure 4C, the comparison displayed higher z-scores for pathways related to viral infection, cell proliferation, survival, and cell movement, highlighting their more active state in processes that are critical for tumor growth and metastasis. This particular finding points out, that the 3D condition of LC4 culture reflects a better model for studying aggressive cancer behavior compared to the 2D culture. Conversely, the HCC core tissue displays higher activation of pathways associated with necrosis (9), sensitivity (2, 3), and apoptosis (10), showing that under in-vivo conditions, a increased stress level is leading to increased cell death and tissue damage. These pathways are significantly downregulated in LC4 cells, implying that the cultured cells are more resistant to stress and apoptotic signals, likely due to the controlled environment of in vitro culture.

Taken together, the LC4 cell line, and particularly under 3D culture conditions, aligns with the parental HCC core in terms of overall pathway activity. While we could show, that LC4 cells exhibit enhanced transcriptional, protein processing, and proliferative activities, reflecting their adaptation to in-vitro conditions, the parental HCC core tissue shows higher activation of tumor-specific and stress-related pathways, such as necrosis and apoptosis.

### LC4 Patient-Derived Cells Retain Immune Evasion and Proliferative Traits but Show Reduced Immune Marker Expression Compared to Parental HCC Core

To explore the adoption and preserved characteristics of the patient-derived LC4 cells in terms of liver specific and immune specific properties, we perform a details comparison using the parental tissue, the isolated cells and the generated cell lines. We aimed to determine whether the LC4 line maintains the diverse cellular components of the parental center region and if certain populations, particularly tumor associated immune cells, as TAMs are diminished during culture establishment. As shown in **Figure 5A** liver- and tumor-associated markers, such as AADAC, CLRN3, GPC3, CD40 and ALB were downregulated in LC4 cells compared to freshly isolated tumor core cells and parental tissues. In contrast, the marker CD47 was consistently expressed, and CD44 appeared to be upregulated in comparison to the parental core region. The persistent expression of CD44 and CD47, linked to stemness and immune evasion, suggests these properties are retained in LC4 cells. In **Figure 5B** a clear reduction in gene expression of immune cell markers for identifying TAMs and MDSCs (e.g., CD68, ARG1, CD163, and S100A8/A9) was detectable in LC4 cell conditions, whereby the markers CD163 and ITGAM were upregulated under 3D culture conditions. The cytometry analysis shown in **Figure 5C** points out, that a minor subset of the LC4 cells (11%) presenting CD68 at very low levels. As shown by immunofluorescence data, represented in **Figure 5D**, LC4 cells retained markers which further supports the result that LC4 cells preserve the proliferative and immune-evasive properties (e.g., Ki67, PDL-1, and CD44) of the parental tumor core. Next, as shown in **Figure 5E and F**, we analyzed MHC class I, class II molecules and Hif1a in the parental samples compared to their cultured counterparts. We found, that while HIF1a and HLA-G were upregulated in expression, while HLA-F was expressed to the same intent as in the core region. MHC class I molecules HLA-A, B and C were expressed to lower amounts, but detectable, while the expression pattern of MHC-class II molecules appeared steady reduced in the LC4 cells.

**Figure 5:**
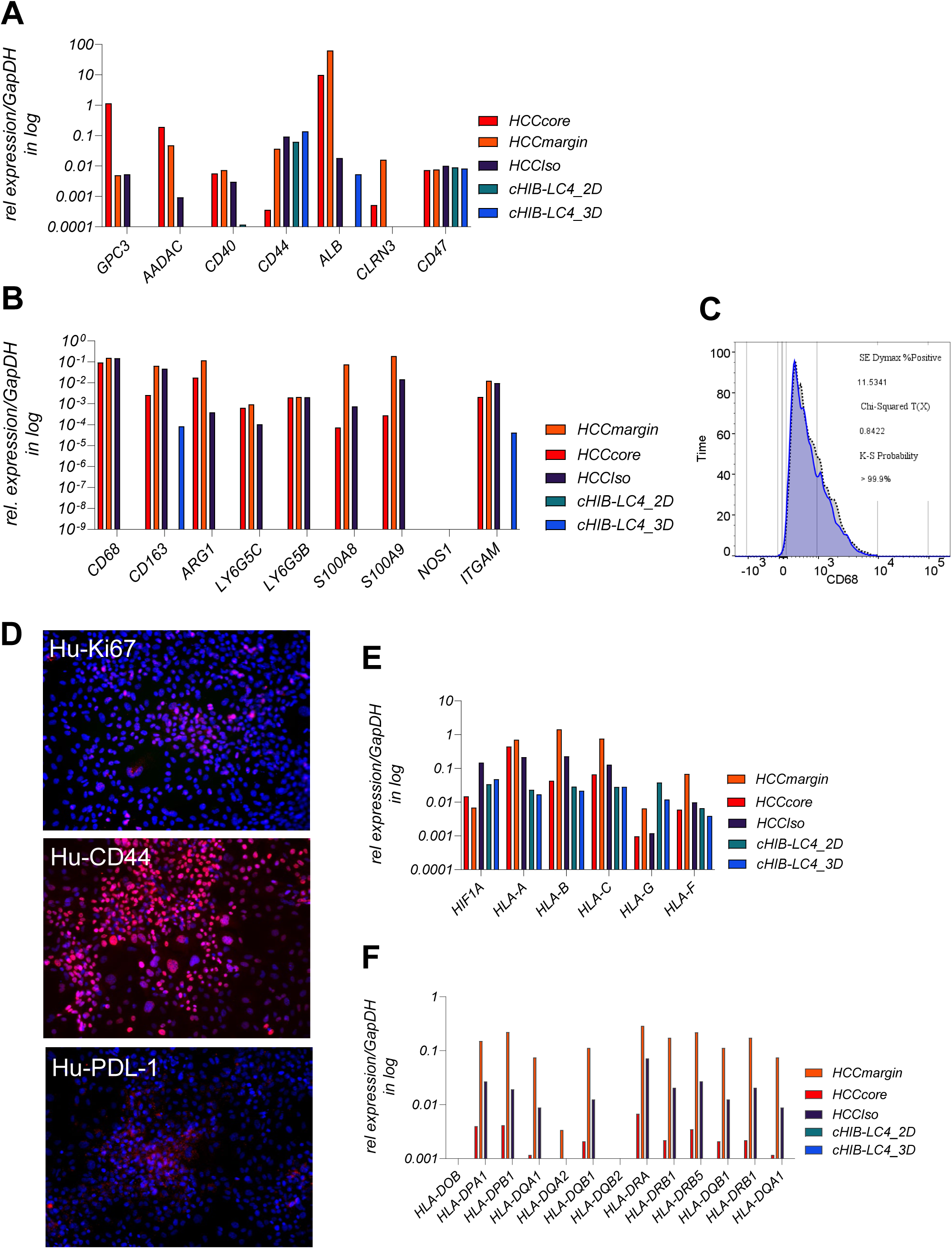
Liver tumor and immune recognition related gene and protein expression profiles of patient-derived LC4 cells compared to the parental specimens. **Figure 5A, B, F and G** displaying mRNA expression levels of selected genes in different cell fractions: HCC core, margin tissue, HCCIso (freshly isolated tumor cells from the core region), LC4_2D (generated cell line in 2D culture conditions), and LC4_3D (generated cell line in 3D culture conditions), normalized against GapDH expression and plotted on log10 scale. In **Figure 5A**, liver and tumor specific genes including GPC3, CD40, CD44, CD47, AADAC, ALB, and CLRN3 were analyzed in the indicated fractions. **Figure 5B** shows the mRNA expression level of genes associated with tumor-associated macrophages (TAMs) and myeloid-derived suppressor cells (MDSCs) in the same samples. The genes analyzed include CD68, ARG1, CD163, ITGAM, LY6G5B, LY6G5C, NOS1, S100A8, and S100A9. This comparison highlights the differences in the expression of immune-related genes under different experimental conditions, showing changes in the tumor microenvironment during cell culture and generating the LC4 cell lines. **Figure 5C** presents flow cytometry analysis for CD68, a marker for macrophages, in LC4 cells. The plot displays the percentage of CD68-positive cells, along with statistical parameters such as the K-S probability, Chi-Squared value, and SE Dynmax %Positive. **Figure 5D** shows immunofluorescence staining of LC4 cells in 2D culture conditions for Ki67, PDL-1, and CD44. The images display the expression of Ki67 (red), PDL-1 (red), and CD44 (red), with nuclei counterstained with DAPI (blue). These staining patterns illustrate cell proliferation (Ki67), immune checkpoint expression (PDL-1), and stemness markers (CD44), offering the visual confirmation of the molecular profiles. In **Figure 5E** and **5F** the differential expression pattern of HIF1A, MHC class I and MHC class II related genes were shown detected in the patient derived tissue regions and cells compared to the generated LC4 cell line.

### LC4 Cells in 3D Culture Exhibit Enhanced Proliferation and Immune Evasion

To better characterize the newly generated, patient-derived LC4 cell line, we conducted a comprehensive differential gene expression (DEG)-based pathway analysis, comparing LC4 cells to healthy hepatocytes. As shown in Figures 6A and 6B, volcano plot visualizations of DEG results laid the foundation for our pathway analysis. For LC4 cells, Figure 6A confirmed findings from Figure 4, showing upregulation of genes related to protein synthesis and signal transduction (e.g., RPL30, SPTAN1), reflecting their adaptation to artificial conditions that promote rapid growth. However, downregulation of genes involved in the extracellular matrix, metabolism, and immune response (e.g., COL3A1, NNMT, C4A, HLA-A) was observed. Figure 6B revealed that 3D-cultured LC4 cells more closely retained key characteristics of the parental tumor core when compared to healthy hepatocytes. In 3D culture, LC4 cells upregulated genes involved in cytoskeletal dynamics (e.g., CORO1B, CCT7) and immune evasion (e.g., PDL1, JAK2), suggesting that 3D conditions better replicate the tumor core’s cellular environment. Interestingly, the downregulation of genes related to immune response and lipid metabolism (e.g., CFH, SCARB1) in 3D culture suggests that some aspects of the tumor microenvironment, including immune modulation, are maintained. In the in-silico z-score-based pathway analysis, presented in Figures 6C and 6D, we compared LC4 cells cultured in 2D and 3D conditions with healthy hepatocytes to further explore pathway activations and inhibitions. As shown in Figure 6C, in LC4_2D culture, pathways such as Neutrophil Degranulation, EIF2 Signaling, and Macrophage Production of Nitric Oxide and Reactive Oxygen Species were upregulated, indicating an inflammatory milieu. In line with findings from Figure 5, MHC class II proteins were downregulated, reflecting immune modulation and stress adaptation, even compared to healthy hepatocytes. Additionally, EIF2 signaling, a key pathway responding to cellular stress, showed high activation. However, pathways related to Eukaryotic Translation Elongation and Protein Ubiquitination were downregulated, suggesting reduced protein synthesis and post-translational modification in 2D culture, diverging from the patterns observed in the parental tumor tissue. In Figure 6D, LC4_3D cultures showed increased activity in pathways related to Neutrophil Degranulation, EIF2 Signaling, and Mitotic Metaphase and Anaphase, indicating an inflammatory environment, enhanced protein synthesis, and increased cell division. The upregulation of mitotic pathways highlights the highly proliferative nature of tumor cells in 3D culture, which better mimics the spatial structure of the parental tumor. However, pathways such as Protein Ubiquitination, Class I MHC Antigen Processing and Presentation, and the RHO GTPase Cycle were reduced, suggesting impaired immune recognition and altered cytoskeletal regulation.

**Figure 6:**
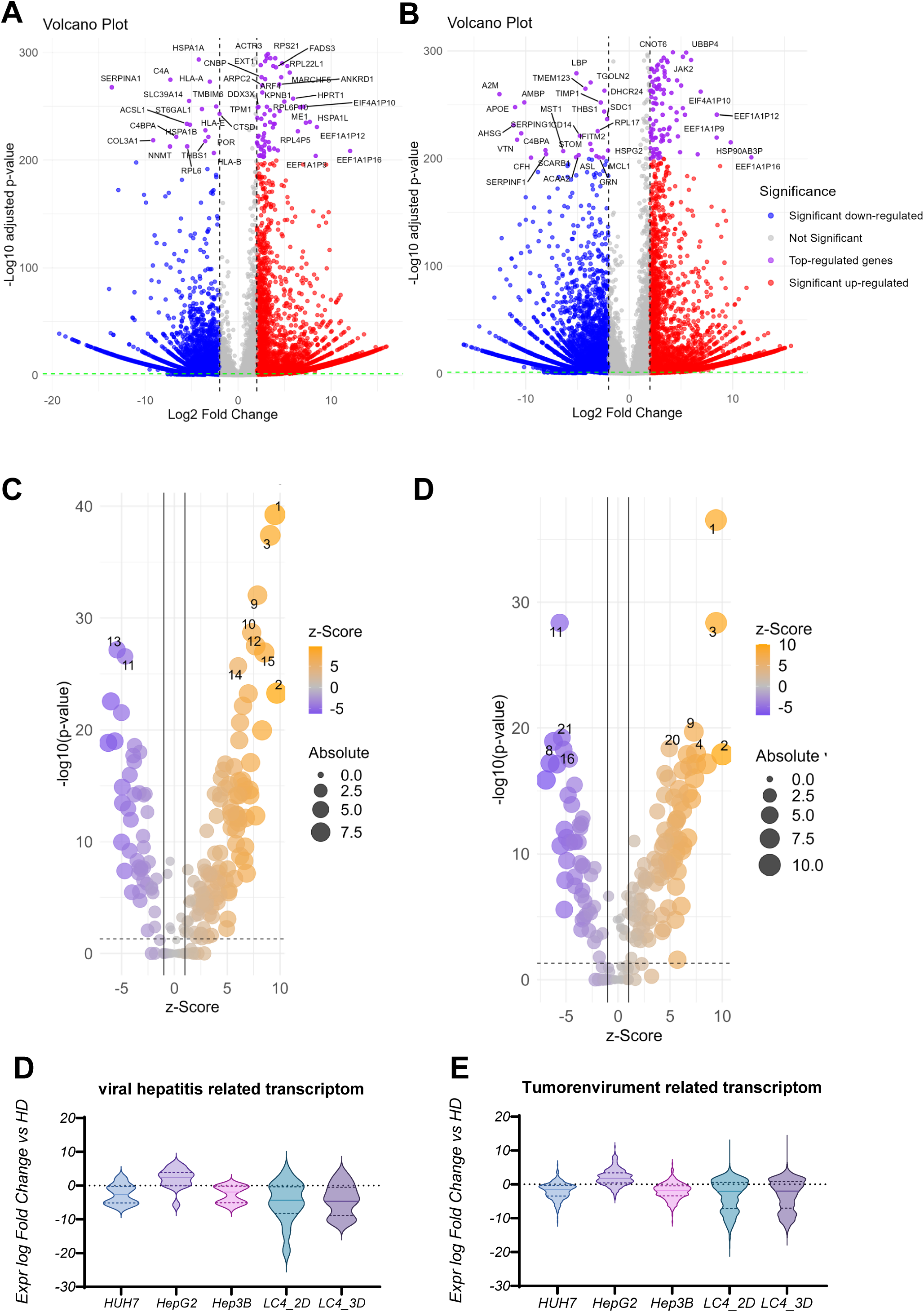
DEG and pathway activation analysis of generated cell line LC4 compared to healthy donors and immortalized cell lines. **Figure 6A and B** showing the differential gene expression between LC4 cells cultured under 2D (LC4_2D; A) and 3D (LC4_3D; B) conditions compared against the expression counts detected in RNASeq of healthy hepatocytes. The volcano plots illustrate the log2 fold change in gene expression (x-axis) versus the -log10 adjusted p-value (y-axis). Each point represents a gene, with the x-axis showing the log2 fold change in expression levels and the y-axis showing the -log10 adjusted p-value. Blue points indicate significantly down-regulated genes (adjusted p-value < 0.05), red points indicate up-regulated considerably genes, and purple points highlight the top-regulated genes based on fold change, with p-values < 200. Grey points represent genes with non-significant changes, defined by adjusted p-value > 0.05 and log2 fold change between 2, -2. In **Figure 6C** the DEG analysis from **Figure 6A** and **B** were used for predictive canonical pathway comparison. The x-axis shows the z-score, indicating the direction and magnitude of expression change, while the y-axis displays the -log10(p-value), representing the statistical significance of the changes. The size of the dots corresponds to the absolute expression levels, and the color gradient indicates the z-score, with orange representing upregulation and purple representing downregulation. Most prominent pathways were numbered and listed in **table 5**. In **Figure 6D and E**, the expression of genes involved in the pathways: viral hepatitis (**D**) and tumor environment (E) were analyzed in fold change against the healthy hepatocytes for commonly used immortalized cell lines Huh7, Hep3B and HepG2 in comparison to generated LC4 cells in 2D and 3D conditions.

### LC4 Cells Exhibit Greater Transcriptomic Changes Compared to Common Liver Cancer Cell Lines, with 3D Culture Retaining More Tumor-Like Characteristics

In the analysis of gene expression related to the viral hepatitis transcriptome, shown in **Figures 6D** and the tumor environment transcriptome, **Figures 6E**, LC4 cells (in both 2D and 3D culture) demonstrate distinct patterns compared to commonly used liver cancer cell lines (HUH7, HepG2, Hep3B). For the viral hepatitis-related transcriptome, HUH7, HepG2, and Hep3B show relatively stable and minimal changes, with expression levels close to those of healthy samples (HD). In contrast, LC4 cells, particularly in 2D culture, display much broader expression changes and stronger downregulation of viral hepatitis-related transcripts compared to the commonly used cell lines. This suggests that LC4 cells, especially in 2D, undergo more significant suppression of viral stress or inflammation-related pathways. LC4_3D exhibits a similar trend but with less pronounced downregulation, suggesting that 3D culture conditions help preserve some viral hepatitis-related characteristics. For the tumor environment-related transcriptome, the commonly used cell lines (HUH7, HepG2, Hep3B) again show minor changes, with expression levels remaining close to those of healthy donors, suggesting limited alteration in tumor environment-related pathways. In contrast, LC4_2D and LC4_3D show broader and more significant downregulation of tumor environment-related transcripts, with LC4_2D exhibiting the strongest reduction. This indicates that LC4 cells, particularly in 2D culture, lose some of the tumor environment-specific features that are present in vivo. LC4_3D, on the other hand, retains more of the tumor environment-related characteristics compared to 2D culture, as indicated by less severe downregulation, making 3D culture a better model for replicating the tumor’s original microenvironment. In direct comparison, LC4 cells demonstrate more significant transcriptomic changes, particularly in 2D culture, compared to the commonly used liver cancer cell lines.

### CYP Enzyme Expression in LC4 Reflects Metabolic Reprogramming in HCC

The CYP enzyme expression, shown in Figure 7, provides key insights into drug metabolism, chemotherapy resistance, and hormonal imbalances in the LC4 model. Enzymes, essential for metabolizing drugs, steroids, and hormones, show significant differences in the expression level of LC4 cells compared to hepatoma cell lines HUH7, HepG2, and Hep3B. In Figure 7A, the downregulation of CYP3A4, CYP3A5, and CYP2C9, which are crucial for metabolizing chemotherapeutics like paclitaxel and docetaxel, are shown. Suggesting higher drug concentrations, increasing toxicity, and diminished activation of prodrugs, contributing to chemotherapy resistance. Figure 7B, reduced CYP19A1 and CYP17A1 expression profiles, impacting estrogen and androgen pathways, were detected in LC4 cells. This imbalance can promote tumor growth and reduce the effectiveness of hormone-based therapies, such as aromatase inhibitors. The disruption in steroid hormone biosynthesis likely affects tumor progression and response to hormone therapies. LC4 cells also uniquely express CYP51A1 and CYP4B1 (Figures 7C, 7D), enzymes linked to cholesterol biosynthesis and xenobiotic metabolism. CYP51A1 suggests alternative metabolic pathways, while CYP4B1 enhances detoxification, potentially increasing chemotherapy resistance by inactivating drugs more effectively. In Figure 7E, CYP downregulation in LC4 cells (CYP2C9, CYP2D6, CYP1A1) mirrors patterns seen in HCC patient samples, especially in patients co-infected with HIV and HCV. This similarity reinforces LC4’s relevance as a preclinical model, accurately reflecting metabolic deficiencies seen in patients. The unique expression of CYP51A1 and CYP4B1 in LC4 cells suggests the model may capture additional metabolic reprogramming, offering insights into drug resistance mechanisms.

**Figure 7:**
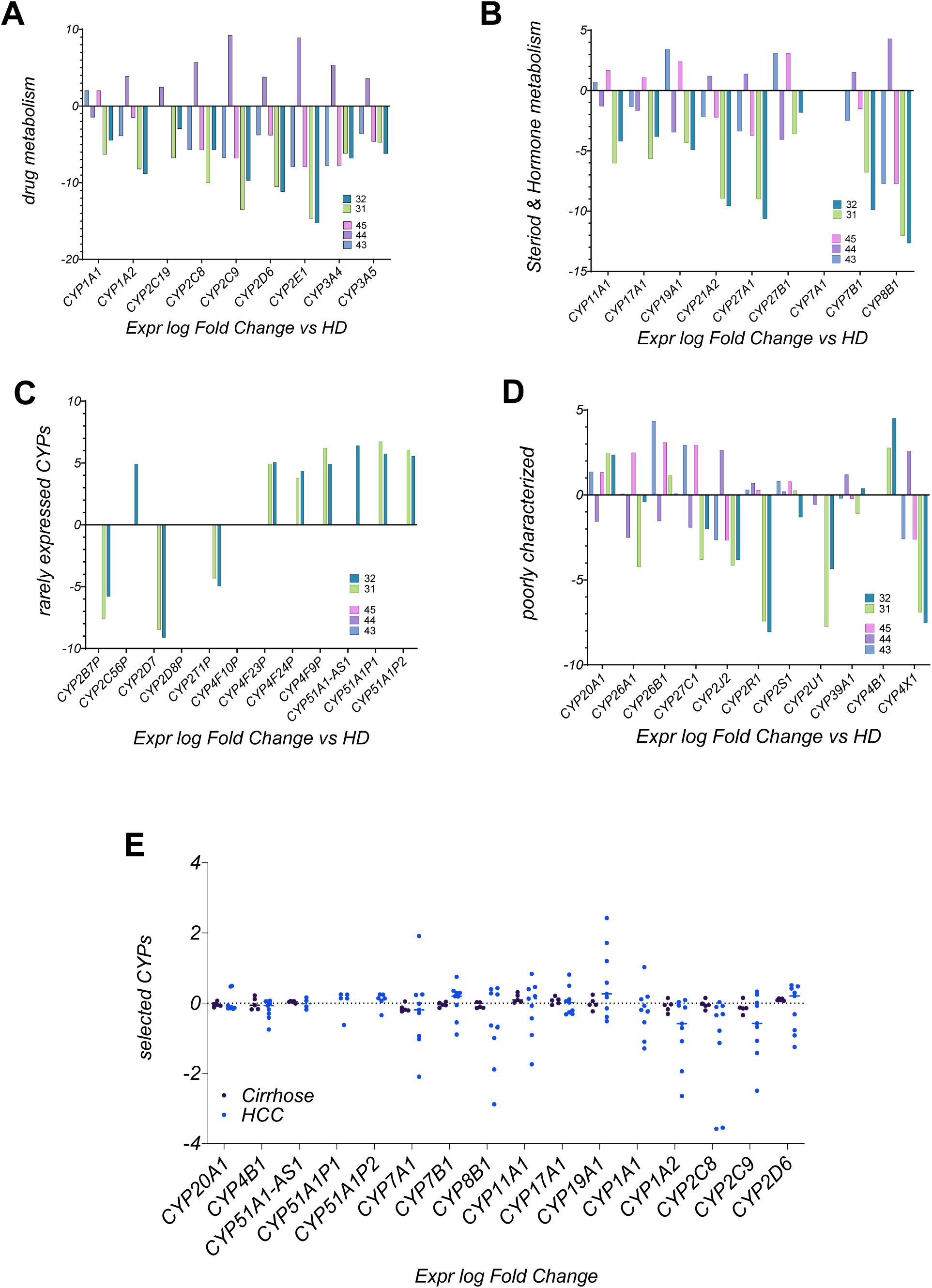
Differential expression analysis of cytochrome P450 (CYP) Enzymes in of patient-derived cell, immortalized cell lines and of global patient Data sets. In the **Figures 7A-B** bar plots visualizing log fold change analysis of CYP enzymes involved in drug metabolism (**A**) and steroid and hormone metabolism (**B**). Rarely expressed and poorly characterized CYPs were displayed in **Figure 7C and D**. The expression profiles were analyzed using healthy donor samples for normalization. The indicated CYP enzymes were analyzed in the conditions LC4 cells in 2D culture (31), and LC4 cells in 3D culture (32), the immortalized cell lines HUH7 = 43, HepG2 = 44, and Hep3B = 45. In Figure 7E a dot plot was used for comparing the expression log fold change of indicated CYP enzymes in a global IPA provided data set of patients with untreated cirrhosis (n = 5) and untreated diagnosed HCC (n = 9).

### Biomarker identification in LC4 cells reveals CDKN2A, IKFZ1 and B2M as consistently regulated

Figure 8 presents an in-depth comparison of biomarker profiles across HCC core, LC4_2D, and LC4_3D cultures, revealing both conserved molecular characteristics and condition-specific adaptations. Figure 8A illustrates the overlapping of investigated biomarkers. In Figure 8B, the biomarker filter process is illustrated. Here, we reduced the markers to a relevant group used for diagnosis and efficacy observations. In this process we identified 52 key biomarkers which were consistently expressed, with particular emphasis on those that could impact therapeutic efficacy and serve as drug targets. Figure 8C categorizes these biomarkers into 14 consistently downregulated and 5 consistently upregulated markers. The heatmap in Supplementary Figure 4A further gives a more detailed picture of the regulated biomarkers. To understand the broader impact of the filtered biomarkers, a global dataset from the IPA platform was employed to compare them with HCCs from various entities. As a result, we identified biomarkers - IKZF1, B2M, and CDKN2A – which showed equal expression profiles along all analyzed specimens.

**Figure 8:**
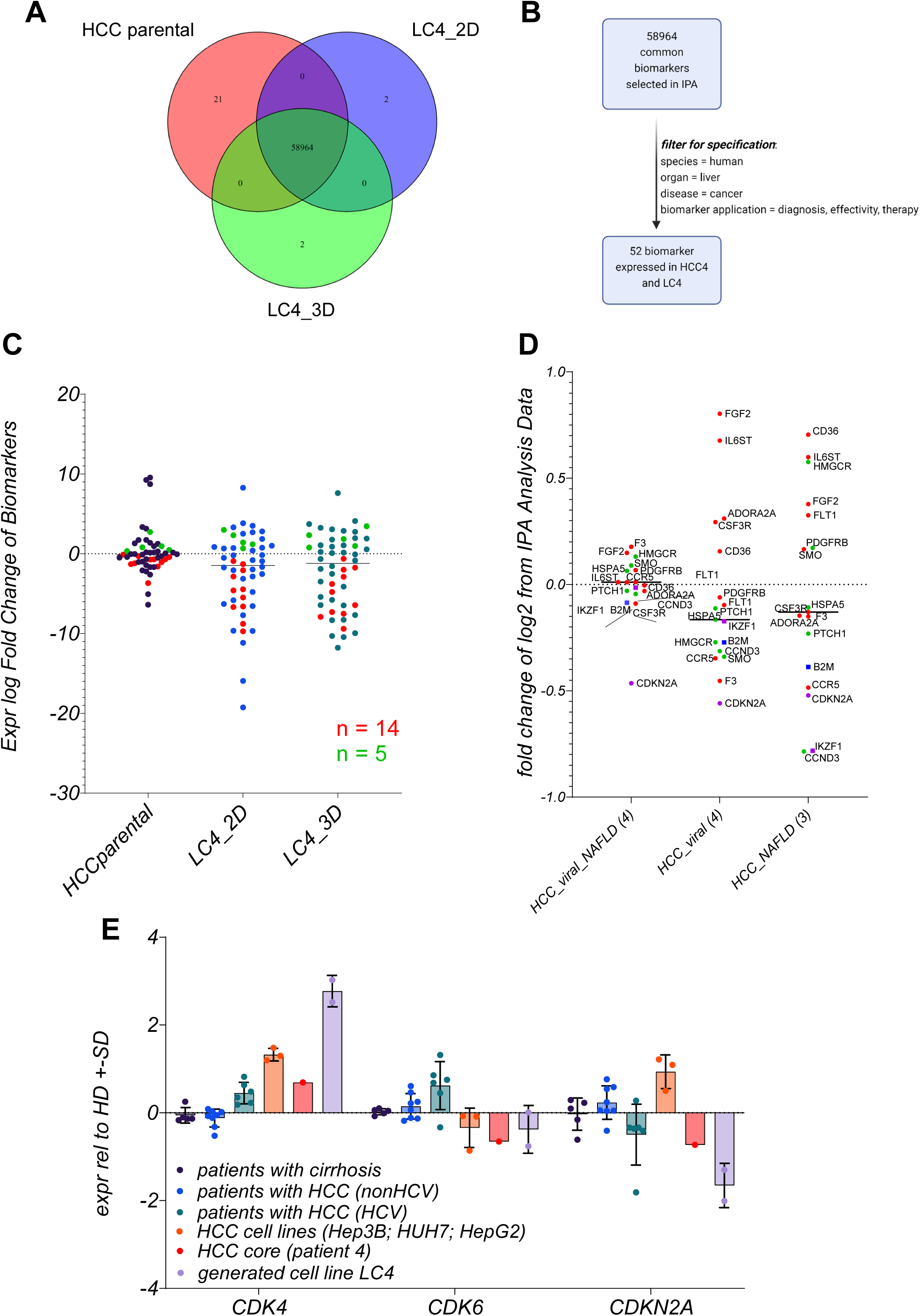
Identification and classification of disease related consistently expressed biomarkers presented in LC4 cells under different conditions. **In Figure 8A**, the Venn diagram illustrating the overlap of biomarkers analysis by IPA biomarker comparison, identified in three experimental conditions: HCC core tumor tissue (HCC core), LC4 cells in 2D culture (LC4_2D), and LC4 cells in 3D culture (LC4_3D). The diagram shows the number of unique and shared biomarkers across these conditions, with a core set of 58,964 common biomarkers identified in all three conditions. In **Figure 8B** the filter progress within the IPA software (Ingenuity Pathway Analysis) was visualized schematically. The selection was filtered for human liver cancer biomarkers relevant to diagnosis, effectiveness, and therapy, resulting in 52 biomarkers expressed in both parental tumor isolated cells and LC4 cells. In **Figure 8C** the scatter dot plot displays the expression log fold change of the selected 52 biomarkers across the three experimental conditions (HCC core, LC4_2D, and LC4_3D). Each dot represents a biomarker, and the distribution illustrates the consistence in the regulation among the different conditions. The green dots indicate consistently upregulated biomarkers (n = 5), and the red dots indicate consistently downregulated biomarkers (n = 14). In the **Figure 8D**, the dot plot analysis shows the fold change of consistently regulated biomarkers from patient 4 derived cells in log2, based on a global IPA analysis data set, comparing relevant specific conditions in HCC patients, by displaying the median of 4 viral/NAFLD patients, 4 viral HCC patients, and 3 NAFLD patients. Here red dots indicating the biomarkers which have been detected to be downregulated in patient 4 derived cells, and green dots indicating such biomarkers which were detected to be upregulated in patient 4 derived cells. Purple and blue squares as well as purple dots identifying the biomarkers which were also consistently regulated in all three conditions of HCCs. In the bar plot displayed **in Figure 8E** the expression log fold change of CDK4, CDK6, and CDKN2A across different patient groups and cell lines compared to healthy donors. The analysis includes samples from patients with cirrhosis (n = 5), patients with HCC (non-HCV; n = 8), patients with HCC (HCV; n = 6,), HCC cell lines (Hep3B and HUH7), the patient 4 isolate (HCC4Iso n = 1), and the generated cell line LC4 (n 02). The standard deviation is represented by the error bars of each column.

As shown in Supplementary Figure 3A, IKZF1 is curial to pathways involved in immune regulation, such as Th2 signaling, B cell receptor signaling, and JAK/STAT pathways. Its downregulation in LC4 cells reflects an immune suppressive milieu, impairing lymphocyte development and contributing to tumor immune evasion. These findings aligns with our previous observations, where LC4 cells displayed reduced immune function and enhanced immune evasion. The Supplementary Figure 3B focuses on B2M, a key component of the MHC Class I complex crucial for antigen presentation. The downregulation of B2M in LC4 is in line with the results presented in Figure 5. The reduced antigen presentation is likely to impair cytotoxic T cell and natural killer cell responses. This downregulation is consistent with the immune-suppressive environment of the parental tissue, particularly in the context of PD-1/PD-L1 signaling.

In the Supplementary Figure 3C, CDKN2A, a tumor suppressor involved in cell cycle regulation, is downregulated in LC4 cells. The loss of CDKN2A function allows activation of the ERK/MAPK and HIF1α signaling pathways, which promotes cell proliferation and survival under hypoxic conditions. The downregulation of CDKN2A mirrors the behavior observed in HCCs, where the loss of cell cycle control facilitates unchecked tumor growth.

### LC4 cells offer a model for CDK6-independent regulation of CDKN2A expression

In Figure 8E the expression level of CDKN2A, CDK4, and CDK6 was compared across different patient groups, including patients with cirrhosis, HCV-related HCC, non-HCV-related HCC, and the parental tissue. LC4 cells display a notable downregulation of CDKN2A accompanied by upregulation of CDK4, similar to the molecular profile of HCV-related HCC patients. This shows that LC4 cells mimic the CDKN2A-CDK4/6 axis of viral HCCs. Interestingly, this expression pattern is distinct from that observed in immortalized cell lines, where CDK4 is upregulated but CDKN2A and CDK6 exhibit varying levels of expression. The consistency of LC4 cells with patient-derived HCC samples highlights their relevance as a model for studying therapeutic responses. The aliment with patient data points to the value of LC4 cells in investigating not only CDK4/6 inhibition but also their interaction with immunotherapies or anti-fibrotic treatments, given the downregulation of B2M and IKZF1.

## Discussion

The present study underscores the complexity of viral-induced HCC by examining the spatial heterogeneity within a single tumor and the usage of LC4 cells derived from that tumor as a pre-clinical model, as the global comparison showed highest similarity to HCC and HCC-related diseases. In this study we offered a complex comparison of different conditions and tissue regions to extract a classification for a newly generated cell line in the setting of HCC research. By comparing the parental tumor core and margin distinct molecular profiles revealed, with the tumor margin showing elevated expression of markers such as CD44 and HIF1A, consistence with stemness, hypoxic adaptation, and metabolic reprogramming at the invasive front. In contrast, the core region exhibited a high proliferation rate, as indicated by Ki67 upregulation, a characteristic mirrored in LC4 cells. These features align with our characterization of LC4 cells, which showed elevated expression of CD44 and Ki67, along with significant cell cycle dysregulation, replicating the dynamics observed in viral-induced HCC [31]. The upregulation of CDK4 and downregulation of CDKN2A in LC4 cells further reflect the aggressive phenotype commonly found in HCC patients [32], making LC4 cells a suitable model for investigating the efficacy of CDK4/6 inhibitors in combination therapies [33]. A critical feature of HCC, particularly in viral infections, is metabolic reprogramming. We observed significant downregulation of CYP enzymes (e.g., CYP3A4, CYP2C8) in LC4 cells, a hallmark of chemotherapy resistance [34]. Reduced expression of these enzymes impairs drug metabolism, leading to higher drug toxicity and diminished activation of prodrugs [35, 36]. These findings are in line with previous results, showing that HepG2 cells, overexpressing CYPs, exert lower resistance to chemotherapy [16]. Interestingly, the upregulation of CYP51A1 in LC4 cells points to an alternative metabolic pathway, involving cholesterol biosynthesis. Cholesterol metabolism has been increasingly implicated in tumor survival and proliferation [37]. The fact that LC4 cells clearly adopt this pathway, and the parental HCC cells were generally characterized as LXR/RXR activating, suggests that inhibitors of cholesterol biosynthesis, such as statins, could be explored as an adjunct therapy to reduce tumor growth, in this cell line. This emphasizes the importance of LC4 cells for investigating the link between metabolic reprogramming and drug resistance.

One of the most significant observations in LC4 cells is their ability to model the immune-suppressive environment found within the parental tumor, mainly through the downregulation of B2M, a critical component of the MHC Class I antigen presentation pathway and the inhibition of IKZF1, a transcription factor involved in lymphocyte recruitment, highlight key mechanisms of immune evasion. The reduction in B2M expression reflects a well-known mechanism of immune evasion, where tumor cells reduce their visibility to cytotoxic T cells, helping them evade immune detection [38]. This is particularly relevant in the context of virus-induced HCC, where chronic immune stimulation from infections drives immune exhaustion and suppression [39]. The downregulation of B2M in LC4 cells mirrors what has been observed in clinical HCC cases, where loss of antigen presentation is associated with resistance to immune checkpoint inhibitors, such as PD-1/PD-L1 therapies [40, 41]. The potential for LC4 cells to model the immune evasion strategies makes them an important tool for investigating combination therapies that could overcome the resistance. For instance, therapies aimed at restoring MHC Class I expression could be combined with checkpoint inhibitors to boost the immune system’s ability to recognize and destroy tumor cells, a strategy that has shown potential in preliminary studies. Additionally, the observed dysregulation of IKZF1 further supports the utility of LC4 cells for studying the effectivity and resistance of immune-modulatory therapies [42]. Given the interplay between chronic viral infection and immune suppression, LC4 cells offer a promising platform to explore novel immunotherapeutic strategies that combine checkpoint blockade with agents to restore T-cell function.

While LC4 cells are a valuable tool for studying HCC biology, they also present limitations, particularly when comparing 2D and 3D culture models. LC4_3D cultures, which better mimic the in vivo tumor microenvironment, offer advantages in studying tumor-stromal interactions, essential for understanding tumor progression and therapeutic responses [43, 44]. The spatial organization captured in 3D cultures makes them more physiologically relevant. However, the limitations of LC4_2D cultures should not be ignored. Although they do not fully replicate tumor-stromal dynamics, 2D cultures remain useful for investigating tumor-intrinsic properties, such as proliferation and cell cycle dysregulation. For instance, the dysregulation of the CDKN2A-CDK4/6 axis in LC4 cells makes the 2D model particularly suited for preclinical testing of CDK4/6 inhibitors, allowing focused analysis of tumor cell behavior in a controlled environment free from stromal influences [10].

In conclusion, this study provides a comprehensive analysis of the spatial heterogeneity within a viral-induced HCC, highlighting the value of LC4 patient-derived cells as a model for investigating tumor progression, immune evasion, and therapeutic resistance. Despite from the classification as HCC-derived cell line by comparing the global analysis, the consistent regulation of key biomarkers such as CDKN2A, IKZF1, and B2M in LC4 cells, and across various specimens, underscores their ability to reflect specific entities of HCC. Furthermore, the insights into chemotherapy resistance and tumor-stroma interactions, particularly in 3D culture systems, offer valuable perspectives for future research and treatment development.

## Aberration

HCC: Hepatocellular carcinoma
MASH: metabolic steatohepatitis
HCV: Hepatitis C virus
HIV: Human immunodeficiency virus
TME: Tumor microenvironment
PD-L1: Programmed death-ligand 1
VEGF: Vascular endothelial growth factor
CTLA-4: Cytotoxic T-lymphocyte-associated protein 4
PDX: Patient-derived xenografts
RNA-seq: RNA sequencing
DEGs: Differentially expressed genes
CDKN2A: Cyclin-dependent kinase inhibitor 2A
CYP: Cytochrome P450
B2M: Beta-2-microglobulin
IKZF1: Ikaros family zinc finger protein 1
EMT: Epithelial-mesenchymal transition
TAM: Tumor-associated macrophages
MDSC: Myeloid-derived suppressor cells
IPA: Ingenuity Pathway Analysis

## Author contributions

JK initiated and supervised the research study; JK, LS and MD designed the experiments; AH and KS supervised the collection of human samples; JK, LS, GR, MG and TV conducted experiments and acquired data; JK, GR, GM analyzed data; JK, LS, SL, SP, MD wrote the manuscript. All authors discussed the data and corrected the manuscript. All authors had access to the study data and had reviewed and approved the final manuscript.

## Acknowledgments

We thank Tobias Gosau and Ursula Mueller for their excellent work. We thank Claudia Dettmer, Eileen Maly, and Karina Börner for their excellent technical assistance. We thank Manfred Jücker for providing immortalized cell lines HUH-7 and Hep3B. We thank Ulrike Protzer and Karthrin Singethan for providing HepG2 cells. We thank Kristoffer Riecken for providing the LeGO-vector which was used for stable transduction of LC4 cells. And we thank Annika Elise Ahrenstorf and Katja Riemann for the assistance and performance for the HLA Typing.

