## Supplementary Methods for "Patient-derived HCC cells recapitulating the transcriptomic landscape of primary HCV-related liver cancer"

***Data Processing and Differential Expression Analysis.*** Raw sequencing reads (n = 2 per sample) were prepared as R input data sets and further processed with R [23]. Input reads were trimmed for adapter sequences and low-quality bases using Trimmomatic [24]. Clean reads were aligned to the human reference genome (GRCh38) using STAR aligner [25]. The resulting BAM files were processed with featureCounts to generate read count matrices [26]. Differential expression analysis was performed using the DESeq2 package in R [27]. Genes with an adjusted p-value (Benjamini-Hochberg correction) of less than 0.05 and a log2 fold change (log2FC) greater than 1 or less than -1 were considered significantly differentially expressed.

***Volcano Plot Generation.*** A volcano plot was generated to visualize the differential expression results using the ggplot2 package in R [28]. The plot displays the log2 fold changes on the x-axis and the -log10 adjusted p-values on the y-axis. Genes with significant upregulation are highlighted in red, while significantly downregulated genes are shown in blue. Genes that did not meet the significance threshold are represented in gray. Genes with p-value > 200 and log fold change <-2 and >2 were classified as top-regulated genes (top-genes) and were entirely listed in the **Supplementary Table 3**.

***Methods for Comparison Analysis in IPA.*** Differentially expressed genes (DEGs) identified from DESeq2 analysis were imported into Ingenuity Pathway Analysis (IPA) software (Qiagen) [29]. Canonical pathway analysis identified pathways significantly enriched. The significance of the association between the dataset and the canonical pathway was measured using the ratio of mapped genes to total genes in the pathway and Fisher's exact test p-values.

***Bubble Plot.*** Following the DEG analysis, z-scores and p-values for DEGs were extracted from IPA with z-scores greater than 5 and p-values greater than 1.3 were visualized. The data were imported into RStudio, where p-values were transformed to -log10 values. A bubble plot was created using ggplot2 to visualize the relationship between z-scores and -log10(p-values) [28]. The size of the bubbles corresponded to statistical significance, and the colour indicated z-score values. Single pathways were numbered in the graphic and listed in detail in **Table 1**.

***Z-score based heatmaps.*** Data were filtered to include only canonical pathways, Biofunctions and diseases or toxicity related pathways with z-scores > 5 and p-values < 0.05. A heatmap was generated using the pheatmap package in R to visualize z-scores of selected pathways [30]. Single pathways were numbered in the graphic and listed in detail in the **Supplementary Table 4 and Table 2 and 3**.

***Correlation Heatmap****.* Following pathway identification in IPA, z-scores and p-values were extracted and imported into RStudio. Pearson correlation coefficients were calculated for z-scores across samples, and the correlation matrix was visualized as a heatmap using the corrplot package in R. The heatmap represented the strength and direction of correlations with a color gradient for correlation coefficients [31].

***Comparative Pathway Analysis Visualization in RStudio.*** Significant pathways related to HCC were selected from IPA, and z-scores and p-values were extracted for visualization in RStudio. Data were imported and transformed to -log10(p-values). A scatter plot was generated using ggplot2, with the x-axis representing pathways and the y-axis representing z-scores. Color differentiation indicated pathway activation and inhibition [28].
