## Supplementary Table 1 for "Patient-derived HCC cells recapitulating the transcriptomic landscape of primary HCV-related liver cancer"

**Supplementary Table 1:** antibodies for processed histological staining

| antigen (clone) | supplier |
|---|---|
| anti- calnexin (C5C9) | Cell Signaling Technology |
| anti- CD3 (CD3-12) | BioRad |
| anti- CD44 (Pa5-114983) | ThermoFischer |
| anti- HCV Core (C7-50) | Abcam |
| anti- HNF4alpha (PA5-18363) | ThermoFischer |
| anti- ki67(OTI93) | Origene |
| anti- PD-L1(IHC411) | GeneTex |
| anti- vimentin (E-5) | Santa Cruz |
| anti-CD68 PG-M1) | Dako |
