## Supplementary Table 2 for "Patient-derived HCC cells recapitulating the transcriptomic landscape of primary HCV-related liver cancer"

**Supplementary Table 2:** TaqMan assays (thermo fischer)

| Gene name | Assay-ID |
|---|---|
| AADAC | Hs00153677_m1 |
| ALB | Hs00609411_m1 |
| CD3G | Hs00173941_m1 |
| CD4 | Hs01058407_m1 |
| CD44 | Hs01075864_m1 |
| CD8A | Hs00233520_m1 |
| CLRN3 | Hs00380707_m1 |
| GapDH | Hs99999905_M1 |
| GZMB | Hs01554355_m1 |
| HIF1A | Hs00936375_m1 |
| IL10 | Hs99999035_m1 |
| MKi67 | Hs04260396_g1 |
| RPL0 | Hs00420895_gH |
| RPL30 | Hs00265497_m1 |
| SLC2A1 / Glut1 | Hs00892681_m1 |
| TNFalpha | Hs99999043_m1 |
| TP53 | HS00931461_M1 |
