## Supplementary Figure 1 for "Patient-derived HCC cells recapitulating the transcriptomic landscape of primary HCV-related liver cancer"

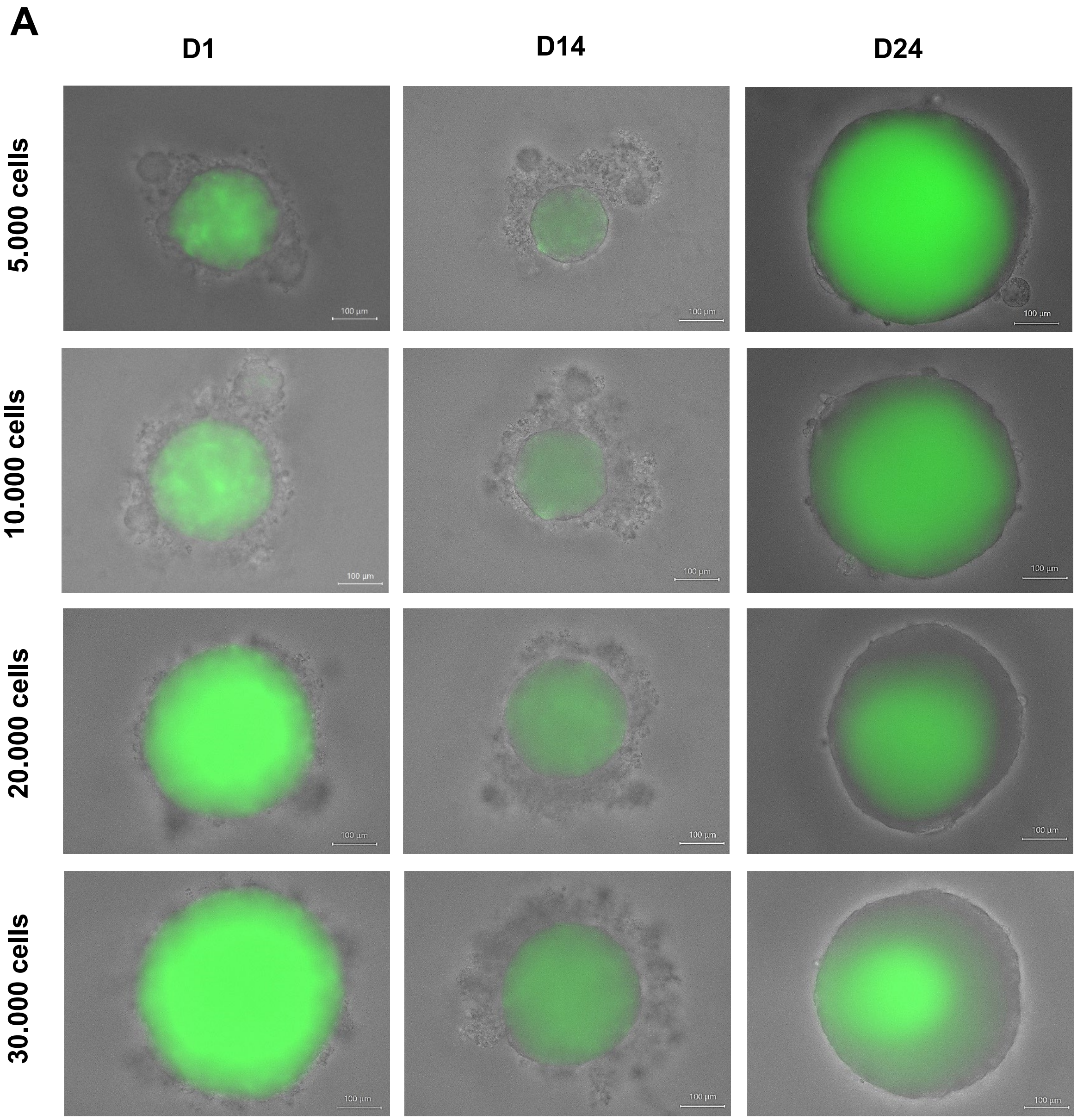

**Supplementary Figure 1: Representative longitudinal Observation of spheroid formation.**

**Supplementary Figure 1** shows the time and cell number dependent formation of LC4 cell stable producing GFP. Cells were seeded on 96-well 3D culture plates with indicated cell numbers. Each well represented one formed spheroid. On indicated timepoints spheroids were captured consistently in the same wells. Visualization of the 3D spheroids was realized using the z-stack function of the fluorescence microscope BZ-9000 from Keyence (Osaka, Japan). Captures were automatically generated with the indicated magnification and consistent exposure for brightfield and GFP. Brightfield and GFP channels were merged for the visualization after preparing a full fucus of the single takes of each channel.
