## Supplementary Figure 2 for "Patient-derived HCC cells recapitulating the transcriptomic landscape of primary HCV-related liver cancer"

| molecule | HCC core | LC4_2D | LC4_3D |
|---|---|---|---|
| ADORA2A | -1.596 | -6.683 | -7.314 |
| AN1A2 | -1.297 | -0.923 | -0.638 |
| B2M | -0.063 | -1.648 | -4.126 |
| CCND3 | 0.482 | 1.414 | 0.776 |
| CCR5 | -0.703 | -4.451 | -7.519 |
| CD36 | -1.195 | -9.727 | -7.912 |
| CDKN2A | -0.73 | -1.30 | -2.01 |
| CSF3R | -0.309 | -8.781 | -9.405 |
| F3 | -0.417 | -2.082 | -2.889 |
| FGF2 | -3.68 | -6.609 | -4.787 |
| FLT1 | -0.107 | -4.596 | -6.444 |
| HMGCR | 0.34 | 2.315 | 2.326 |
| HSPA5 | 0.947 | 1.181 | 0.964 |
| IKZF1 | -0.035 | -7.701 | -8.324 |
| IL6ST | -1.05 | -2.86 | -1.74 |
| PDGFRB | -0.985 | -9.65 | -8.276 |
| PTCH1 | 2.73 | 0.67 | 1.85 |
| RUN13 | -0.691 | -5.483 | -3.791 |
| SMO | 0.81 | 2.99 | 3.44 |

**Supplementary Figure 2: Detailed Heatmap of consistent Biomarker.**

In the **Supplementary Figure 2,** consistently regulated biomarkers, extracted and filtered in the IPA platform were displayed in a heatmap with expression levels indicated and across the three analyzed conditions, as mentioned in the header of the heatmap. Red color indicates downregulation and blue color indicates upregulation, therefore green gene names were displayed for consistent upregulated genes, while red gene names were displayed for consistent downregulated genes, over conditions.
