## Supplementary Figure 3 for "Patient-derived HCC cells recapitulating the transcriptomic landscape of primary HCV-related liver cancer"

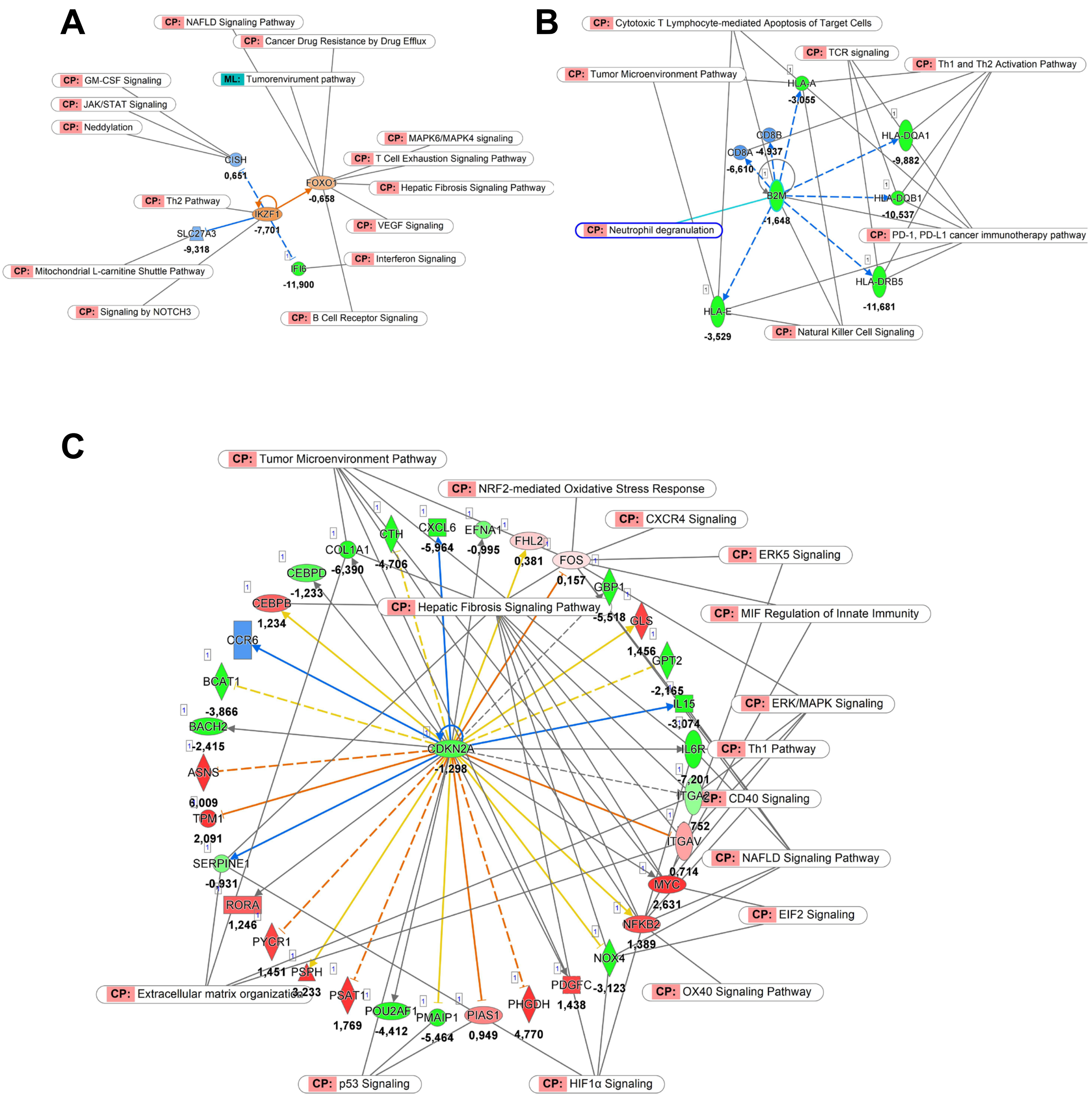

**Supplementary Figure 3: Pathways maps for consistently expressed biomarkers presented in LC4 cells with gene expression and predicate activity.**

In **Supplementary Figure 3A,** the regulatory network diagram illustrates the relationships between the consistently regulated biomarker IKZF1 and its associated canonical pathways (CP) based on IPA (Ingenuity Pathway Analysis). The graph highlights estimated inactivating (blue arrows) and activating (orange arrows) interactions of IKZF1 and downregulated genes in green derived from differential expression analysis in LC4 cells under 2D conditions. The identified pathways are involved in immune responses, cell signaling, and the tumor microenvironment, including Th1 Pathway, T Cell Exhaustion Signaling, and MAPK Signaling, emphasizing the role of IKZF1 in modulating these critical biological processes. In **B** the regulatory network diagram illustrates the relationships between consistently regulated biomarker B2M and its canonical pathways (CP) based on IPA analysis. The graph shows estimated inactivating interactions (blue arrows) of B2M and downregulated genes in green based on the DEG and downstream analysis in IPA of LC4 cells in 2D conditions, identifying pathways being involved in immune responses, cell signaling, and the tumor microenvironment, including TCR signaling, natural killer cell signaling, and PD-1/PD-L1 cancer immunotherapy pathways. In **C** the regulatory network diagram for the consistently regulated biomarker CDKN2A and its canonical pathways (CP) based on IPA analysis is shown. Upregulated genes (red), downregulated genes (green) as well as estimated inactivating (blue arrows) and activating (orange arrows) interactions of CDKN2A are displayed according to the IPA based DEG and downstream analysis in LC4 cells in 2D conditions. The network identifies pathways involved in cell cycle regulation, tumor progression, and the tumor microenvironment, such as the Tumor

Microenvironment Pathway, p53 Signaling, and Hepatic Fibrosis Signaling, underlining CDKN2A's significant impact on these pathways in liver cancer.
